# Prediction of glyphosate resistance level based on *EPSPS* gene copy number in *Kochia scoparia*

**DOI:** 10.1101/047878

**Authors:** Todd A. Gaines, Abigail L. Barker, Eric L. Patterson, Philip Westra, Eric P. Westra, Robert G. Wilson, Andrew R. Kniss

## Abstract

Glyphosate-resistant (GR) *Kochia scoparia* has evolved in dryland chemical fallow systems throughout North America and the mechanism involves 5-enolpyruvylshikimate-3-phosphate synthase (*EPSPS*) gene duplication. Sugarbeet fields in four states were surveyed for *K. scoparia* in 2013 and tested for glyphosate-resistance level and *EPSPS* gene copy number. Glyphosate resistance was confirmed in *K. scoparia* populations collected from sugarbeet fields in Colorado, Wyoming, and Nebraska. The GR samples all had increased *EPSPS* gene copy number, with median population values up to 11. An empirical model was developed to estimate the level of glyphosate-resistance in *K. scoparia* based on *EPSPS* gene copy number. The results suggested that glyphosate susceptibility can be accurately diagnosed using *EPSPS* gene copy number, and further increases in *EPSPS* gene copy number could increase resistance levels up to 8-fold relative to susceptible *K. scoparia*. These trends suggest that continued glyphosate selection pressure is selecting for higher *EPSPS* copy number and higher resistance levels in *K. scoparia*. By including multiple *K. scoparia* samples lacking *EPSPS* gene duplication, our empirical model provides a more realistic estimate of fold-resistance due to *EPSPS* gene copy number compared to methods that do not account for normal variation of herbicide response in susceptible biotypes.

## Introduction

Adoption of glyphosate-resistant (GR) sugarbeet systems can result in improved weed control and reduced sugarbeet injury compared to conventional sugarbeet systems [1]. Glyphosate can provide weed control similar to or greater than conventional weed control programs consisting of three applications of desmedipham, phenmedipham, triflusulfuron, and clopyralid [2]. Prior to commercial introduction, net economic return was predicted to be significantly greater for GR sugarbeet systems compared to conventional sugarbeet due to reduced crop injury and better weed control [1]. GR sugarbeets were commercially introduced in 2007. By 2009, more than 85% of US sugarbeet hectares were seeded with GR cultivars, with remaining areas seeded with conventional cultivars that had resistance to specific pests or diseases that were not commercially available with the GR trait [3]. Sugarbeet growers have significantly reduced tillage and increased net economic return since adoption of GR sugarbeet [4].

*Kochia scoparia* is a competitive weed that can cause substantial yield loss, and is particularly a problem weed in sugarbeet [5, 6]. *Kochia scoparia* is a C4 summer annual broadleaf weed that can germinate and emerge early in the growing season and is tolerant to heat, drought, and saline conditions [5]. *Kochia scoparia* has protogynous flowers in which the stigmas usually emerge one week before pollen is shed and are receptive to foreign pollen which can promote outcrossing between plants in close proximity [7]. It also produces copious amounts of pollen for extended periods of time, which is generally an indication that the species is naturally highly outcrossing [5]. *Kochia scoparia* stem breakage at the soil surface during senescence allows for a tumbling seed dispersal mechanism that can contribute to high rates of spread in the western US [8]. In Wyoming, *K. scoparia* densities as low as 0.2 plants m^-1^ of crop row reduced sugarbeet root yield by 18% [9]. The outcrossing nature of *K. scoparia* combined with prolific seed production results in genetically diverse populations that facilitate the evolution of herbicide-resistance mechanisms [10].

GR *K. scoparia* was first identified in Kansas [11] and has now been identified in multiple Great Plains States including Colorado, South Dakota, North Dakota [12], Montana [13], Nebraska [14], and in the Canadian provinces of Alberta [15] and Saskatchewan and Manitoba [16]. Most of the reported GR *K. scoparia* populations appear to have evolved in reduced‐ or no-till chemical fallow systems [17], where glyphosate is used as the primary weed control practice during fallow periods. Glyphosate resistance in *K. scoparia* has been shown to be due to gene duplication in which resistant plants contain 3 to 10 times more functional copies of the gene encoding 5-enolpyruvylshikimate-3-phosphate synthase (*EPSPS*) [12]. These extra gene copies result in overproduction of the EPSPS enzyme, which is the target enzyme inhibited by glyphosate. High-coverage sequencing analysis of *EPSPS* transcripts from glyphosate-resistant *K. scoparia* revealed the absence of any known resistance-conferring non-synonymous mutations [12], further demonstrating that increased EPSPS protein quantity due to *EPSPS* gene duplication and increased transcription confers glyphosate-resistance in *K. scoparia*. Increased *EPSPS* gene copy number and expression has been shown to be a mechanism for glyphosate-resistance in *K. scoparia* collected from Kansas [18]; Colorado, North Dakota, and South Dakota [12]; and Montana [19]. Detection of *EPSPS* genes on distal ends of homologous chromosomes suggests that increase in *EPSPS* gene copies in GR *K. scoparia* occurred as a result of unequal crossover during meiosis resulting in tandem gene duplication [20]. The extra *EPSPS* copies are stably inherited in *K. scoparia*, consistent with the cytogenetic observation that the extra *EPSPS* copies are located at a single locus [20]. Analysis of *EPSPS* gene copy number and resistance level in *K. scoparia* populations from Kansas suggest that there has been a progressive increase in *EPSPS* gene copies and level of glyphosate resistance over time from 2007 to 2012 [20]. In two GR *K. scoparia* populations from Kansas, *EPSPS* gene copy number was correlated to resistance level, such that within a resistant *K. scoparia* population, individuals with higher *EPSPS* copy number displayed less injury symptoms compared to individuals with lower *EPSPS* copy number [18].

Widespread adoption of GR sugarbeet systems in the US has resulted in significant glyphosate selection pressure, and increasingly sugarbeet growers are reporting reduced *K. scoparia* control with glyphosate. Therefore, the objectives of this study were to a) confirm whether glyphosate resistance was present in *K. scoparia* collected from sugarbeet fields; b) determine whether GR *K. scoparia* from sugarbeet fields has the same mechanism of resistance (increased *EPSPS* gene copy number) as previously identified in dryland fallow systems; c) quantify the effect of *EPSPS* copy number on whole-plant response to glyphosate across numerous (30) *K. scoparia* populations collected from sugarbeet fields from Colorado, Wyoming, Nebraska, and Montana; and d) develop a model which can be used to estimate the level of glyphosate resistance in *K. scoparia* based on *EPSPS* gene copy number without the need for time consuming greenhouse bioassays.

## Materials and Methods

### Plant Material

In the autumn of 2013, 65 sugarbeet fields from Colorado, Wyoming, Nebraska, and Montana were surveyed by Western Sugar Cooperative agricultural staff for surviving *K. scoparia* plants. Seed was collected from maturing *K. scoparia* plants within the sugarbeet fields, or in some cases, along field margins. GPS coordinates were recorded for each sampling site. Seed was stripped by hand from multiple branches of each *K. scoparia* plant and placed in a plastic bag. Each seed sample (accession) represented one to five individuals, and were therefore not necessarily representative of the entire *K. scoparia* population from a given field. This survey was biased for surviving *K. scoparia* plants from fields where glyphosate was used. Seed from each site was sent to the Panhandle Research & Extension Center in Scottsbluff, Nebraska for whole-plant dose response bioassay. Once received, all accessions were air dried and cleaned before use in the dose response studies.

### Greenhouse bioassay

Each *K. scoparia* accession was screened for susceptibility to glyphosate. Approximately 15 to 20 seeds were planted in 10 cm × 10 cm plastic pots filled with a 50:50 by weight mixture of field soil and commercial potting mix. After planting, pots were placed in a greenhouse where air temperature was maintained at 27 C and pots were watered several times per day with an automated sprinkler system. *Kochia scoparia* emerged approximately three days after planting, and the pots were thinned to three plants per pot shortly after emergence.

When *K. scoparia* averaged 10 cm in height, each accession was treated with five rates of glyphosate (Roundup PowerMAX, Monsanto Company) with ammonium sulfate (2% w/v) and nonionic surfactant (0.25% v/v). A nontreated control was also included for each accession so that glyphosate was applied at 0, 435, 870, 1740, and 3480, and 5050 g ae ha^-1^. Each glyphosate rate by accession interaction was replicated six times for a total of 36 pots per accession, with three individual plants in each pot. A total of 65 *K. scoparia* accessions were included in the greenhouse bioassay. Herbicide treatments were applied in a CO_2_-pressurized moving-nozzle spray chamber calibrated to deliver 224 L ha^-1^ spray solution.

*Kochia scoparia* injury in each pot was evaluated visually 14 days after treatment on a scale from 0 to 100 where 0 represented no injury and 100 represented death of all plants in the pot. For analysis, injury evaluations were converted to a binomial response of alive (<99% injury) or dead (≥99% injury). A two-parameter log-logistic model (Equation 1) appropriate for binomial data (alive vs dead) was used to estimate the glyphosate dose causing 50% mortality (LD_50_) for each kochia accession. The log-logistic model is of the form:

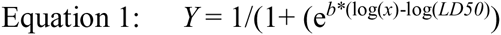

Where *Y* is the probability of survival; *x* is the glyphosate dose in *g* ae ha^-1^; *LD50* is the dose required to cause 50% mortality; and *b* is the slope of the curve at the LD_50_.

### EPSPS gene copy number assay

Based on the results of the greenhouse bioassay, a sub-set of 40 *K. scoparia* accessions exhibiting a range of whole-plant resistance levels were assayed for *EPSPS* copy number. To determine *EPSPS* copy number in these *K. scoparia* accessions, genomic DNA was extracted from individual plants using the DNeasy Plant Mini Kit (Qiagen). Young leaf tissue (100 mg) was sampled from 6-12 plants for each accession when the plants reached 7-10 cm in height. Samples were disrupted in 2 mL tubes using the TissueLyser II (Qiagen). The extraction proceeded using the standard DNeasy Plant Mini Kit protocol. Genomic DNA was eluted in 50 μL of 37 C HPLC water. The concentration and quality of the gDNA were determined using a NanoDrop 1000.

*EPSPS* copy number was estimated using quantitative PCR (qPCR) on the genomic DNA with previously reported primers [12]. Acetolactate synthase (*ALS*) was used as a normalization gene because *ALS* copy number has been shown to not vary among *K. scoparia* individuals [12]. Each reaction contained 12.5 μL of PerfeCTa SYBR^®^ green Super Mix (Quanta Biosciences), 1 μL of the forward and reverse primers [10 μM final concentration], and 5 ng gDNA in a total volume of 25 μL. A BioRad CFX Connect Real-Time System was used for all qPCR. The temperature profile was as follows: 3 min at 95 C followed by 40 rounds of 95 C for 30 sec, 60 C for 30 sec, and 72 C for 30 sec, with a fluorescence reading taken after each round. A melt curve from 65-95 C in 0.5 C increments was performed with a fluorescence reading after each increment to determine the number of PCR products formed in each reaction. Only single PCR products were observed in melt-curve analysis from *EPSPS* and *ALS* primers as expected in all samples, indicating PCR amplification occurred for only the intended genes. The cycle was recorded for each sample at which the fluorescence reading crossed a threshold (C_t_) indicating exponential increase, and relative *EPSPS* gene copy number was calculated using the comparative C_t_ method[21] as 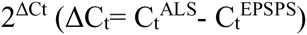 [12]. For each *K. scoparia* accession, 6 to 12 samples were measured for relative *EPSPS* copy number, and the mean, standard error of the mean, and median *EPSPS* copy number were calculated for each accession.

### Effect of *EPSPS*copy number on whole-plant response

A subset of 30 *K. scoparia* accessions were used to quantify the effect of *EPSPS* copy number on whole-plant response to glyphosate. Of the 40 *K. scoparia* accessions analyzed for *EPSPS* copy number, 10 accessions were removed from this analysis due to large standard errors around the LD_50_ estimate from the greenhouse bioassay (S1 Fig). For readers interested in the impact of this decision on the model, results with and without these 10 accessions have been provided (S2 and S3 Figs). The LD_50_ from the greenhouse bioassay was regressed against the median *EPSPS* copy number for each accession. Because the dose response study was conducted at the accession level (that is, multiple individuals from each accession were used as experimental units), the median *EPSPS* value was used to quantify the relationship between gene copy number and whole-plant resistance. The mean copy number could be skewed significantly by a few (or even one) high copy number individuals within a population, and therefore, overestimate the number of high copy number plants for an accession.

We did not know *a priori* which type of regression model was appropriate to quantify the relationship between *EPSPS* gene copy number and whole-plant glyphosate resistance. Based on a preliminary visual evaluation, three different models were fit to the data; a three-parameter rectangular hyperbolic model, a two-parameter rectangular hyperbolic model, and a simple linear model. The three-parameter model was of the form:

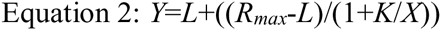

Where *Y* is the LD_50_ from the greenhouse bioassay; *X* is the *EPSPS* copy number; *R*_*max*_ is an upper asymptote, or the maximum theoretical level of resistance at very high values of *X*; *L* is the estimated LD_50_ when *X*=0; and *K* is the value of *X* that results in *Y* halfway between *L* and *R*_*max*_. The two-parameter nonlinear model was the same, but with *L*=0; setting *L*=0 reduced the equation to the Michaelis-Menten model. The linear model used was a simple linear regression. After all three models were fit to the data, Akaike information criterion (AIC) was used to determine which model provided the best fit to the data.

## Results

### Greenhouse bioassay

Of the 65 *K. scoparia* accessions included in the bioassay, 12 had large standard errors associated with the LD_50_ estimates. The LD_50_ results with and without these accessions are provided in S1 Fig. Estimated LD_50_ values ranged from less than 500 to nearly 4000 g ae ha^-1^ (Fig 1A). A majority of *K. scoparia* accessions showed a susceptible response (median LD_50_ for all accessions tested was 906 g ae ha^-1^), even though this survey was biased toward *K. scoparia* plants that were likely to have survived glyphosate application(s). Although the median LD_50_ of 906 g ae ha^-1^ is greater than the standard field use rate of 840 g ae ha^-1^, this is not an indication of a high level of resistance. Previous studies have shown that up to 53-fold more glyphosate is required to control the same *Chenopodium album* biotype in the greenhouse compared to the field [22]. Conversely, greater glyphosate efficacy has been observed in an outdoor environment compared to greenhouse conditions in *K. scoparia*, although both indoor and outdoor environments used artificial growth media rather than field soil [18]. Soil microorganisms present in field soil (but presumably absent from commercial potting media) have been shown to influence glyphosate efficacy in dose response studies [23](Schafer et al. 2012). The LD_50_ (and resulting R:S ratio) from greenhouse and field studies have been recently shown to vary in response to a variety of environmental factors when screening GR *K. scoparia* [24]. Because the LD_50_ can be influenced by a variety of factors, the LD_50_ values in our study (or any greenhouse study) are not an absolute indicator of resistance, but rather a relative measure used to compare accessions within the study for glyphosate response.

**Fig 1.**
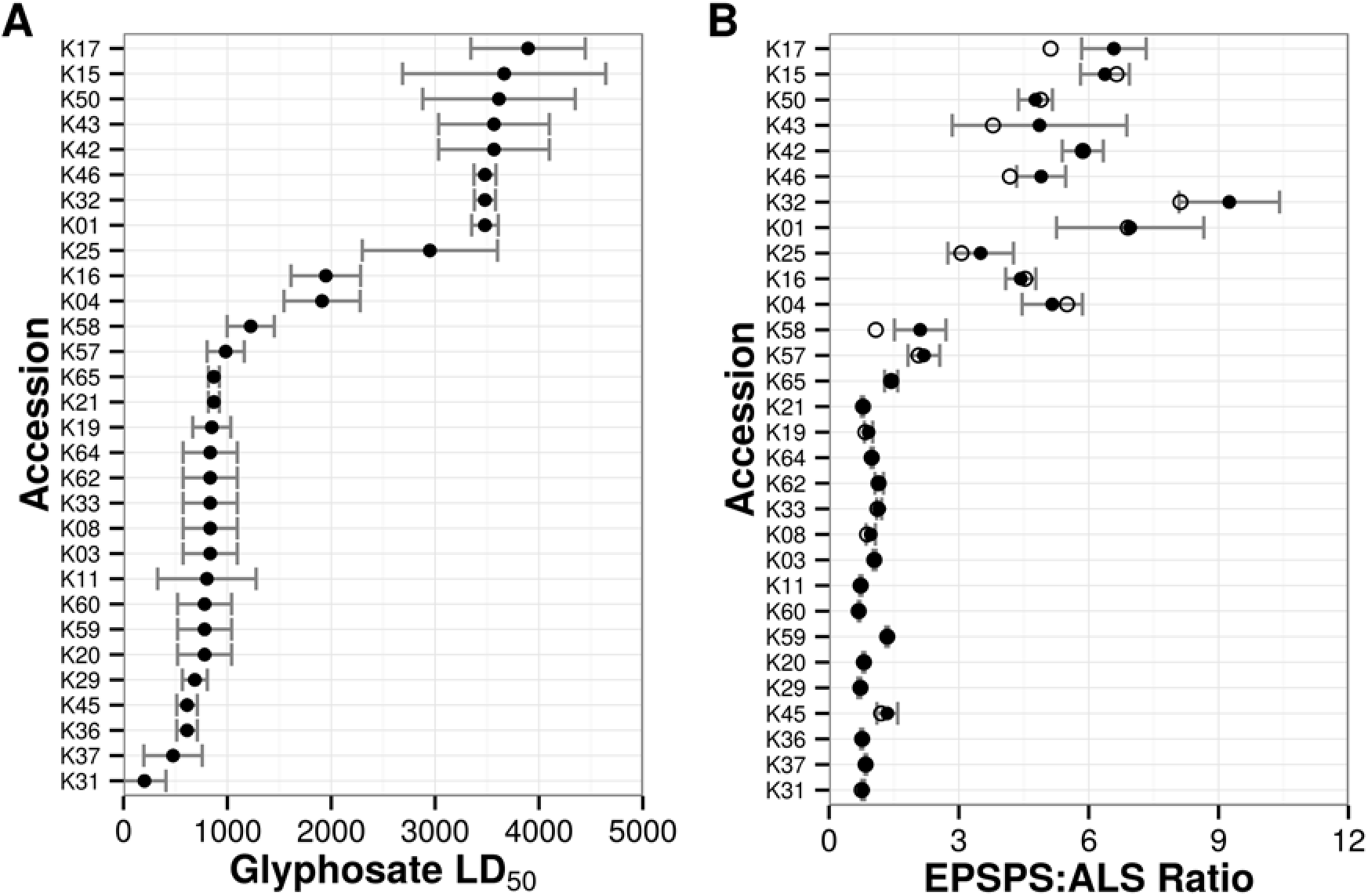
Glyphosate resistance level (A) and *EPSPS* gene copy numbers (B) for 30 *K. scoparia* accessions. Filled circles represent mean values for each accession; bars represent standard error of the mean; open circles represent median *EPSPS* gene copy number for each accession.

### EPSPSgene copy number

Mean *EPSPS* copy numbers in the 30 accessions assessed for effect of *EPSPS* copy number on whole-plant response ranged from 0.7 to 10.2 (Fig 1B), and *K. scoparia* accessions with increased *EPSPS* copy number were identified in Wyoming, Nebraska, and Colorado (Fig 2). Since the accessions were still presumed to be segregating for the GR trait, the median *EPSPS* copy number may provide a more accurate estimate of gene copy level as it relates to resistance level for the accession. Median *EPSPS* copy numbers ranged from 0.7 to 11.3 depending on *K. scoparia* accession. The median *EPSPS* copy number was less than the mean for nearly all accessions where a notable difference was present, indicating that a few individual plants had much higher *EPSPS* copy numbers compared to the majority of plants within that accession.

**Fig 2.**
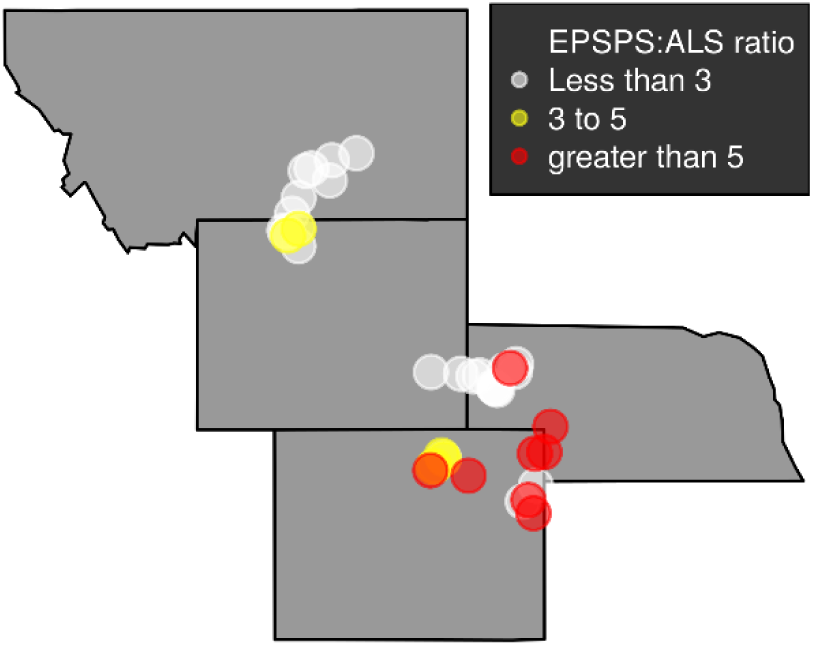
Collection locations of 40 *K. scoparia* accessions for which *EPSPS* copy numbers were quantified.

### Effect of EPSPScopy number on whole-plant response

A notable increase in LD_50_ was apparent for accessions with median *EPSPS* value of >2.5 (Figs 1A and 1B). Based on AIC, the 2 parameter Michaelis-Menten model fit was chosen as the best model (of those tested) to describe the relationship between resistance level and *EPSPS* gene copy number. This suggests there is a ‘plateau’ with respect to resistance level; that is, additional gene copies provide limited increase in resistance level once a certain threshold has been reached. To ensure that the ‘trimmed’ data set did not have a major impact on the empirical relationship, the same models were fit to the full data set of 40 accessions, which included LD_50_ estimates with high standard errors. The results were similar (S3 Fig).

*R*_*max*_, or the theoretical maximum LD_50_ when the number of *EPSPS* gene copies is very large, was 7486 g ae ha^-1^ (Fig 3). The *R*_*max*_ parameter, as an estimate of LD_50_, is not very useful in absolute terms. The LD_50_ value depends heavily on the environmental conditions during the bioassay [18, 25], and can vary significantly from one experimental run to the next even when using the same genotypes and experimental design. However, the LD_50_ ratio between resistant and susceptible biotypes tends to remain relatively more stable than other responses such as GR50 calculated from dry weight [26]. A standardized estimate of resistance level can therefore be calculated by dividing the *R*_*max*_ parameter by the model estimate of LD_50_ for a plant with a single *EPSPS* gene. Practically speaking, this gives an estimate of the “fold” resistance expected due to increased *EPSPS* gene copies. The estimated LD_50_ for a *K. scoparia* individual with a single copy of the *EPSPS* gene in our study was 897 g ae ha^-1^. Our analysis suggests that increasing *EPSPS* gene copies could potentially increase glyphosate resistance level by a maximum of about 8.3-fold (7486 / 897).

**Fig 3.**
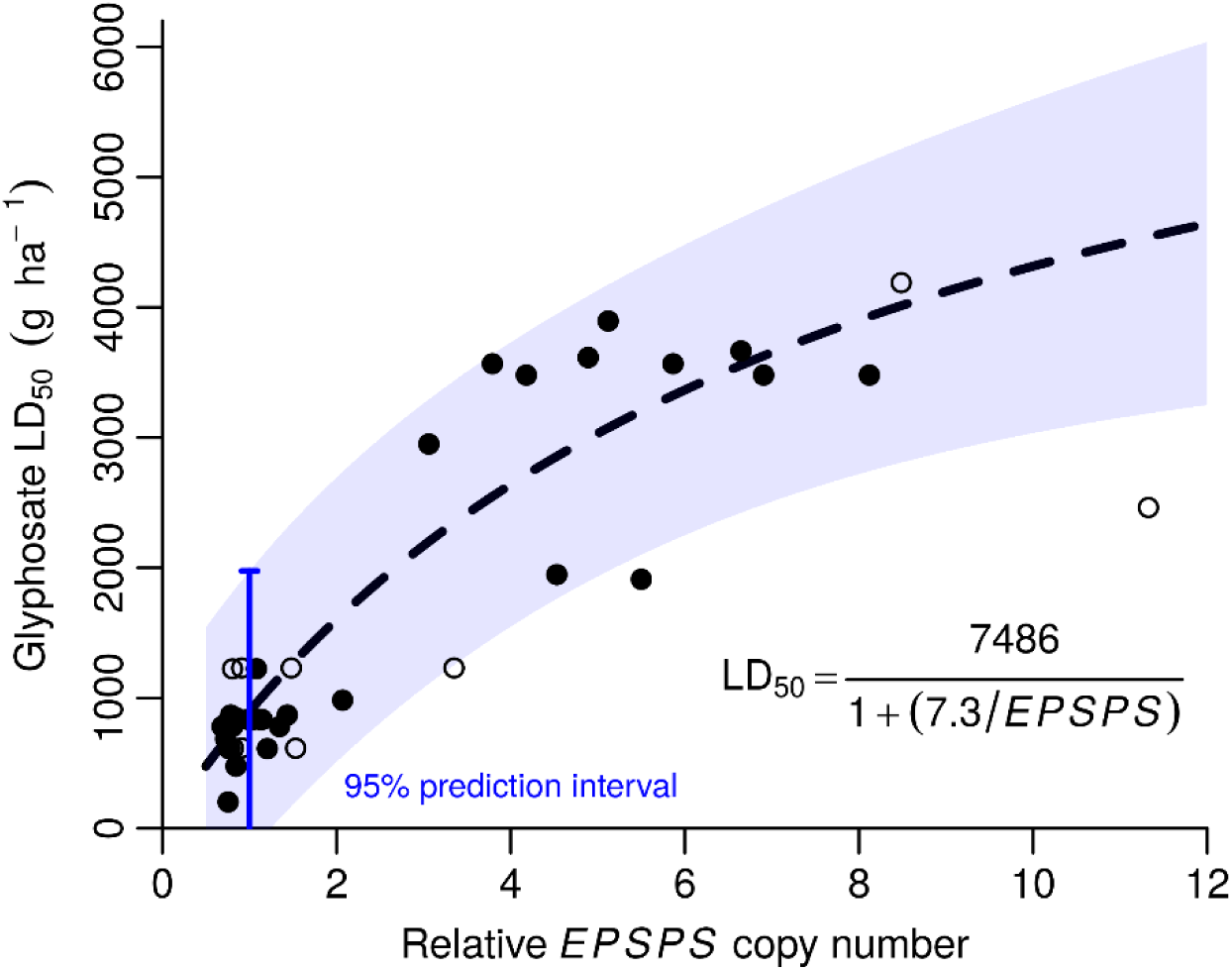
Glyphosate resistance level as influenced by *EPSPS* gene copy numbers in *K. scoparia.* Dotted line represents the fitted model for the 2 parameter Michaelis-Menten (Equation 1). Shaded blue area represents the 95% prediction interval around the fitted line. Filled circles represent *K. scoparia* accessions that were used in the model-fitting (N=30), while open circles represent accessions that were not used in the model fitting procedure (N=10) due to LD_50_ estimates having large standard errors.

The estimated maximum level of glyphosate resistance seems to be in line with previous work on glyphosate resistance, which report 4‐ to 11-fold resistance in glyphosate-resistant *K. scoparia* accessions compared to a single susceptible accession [13, 15, 18]. A generalized version of our model (Equation 3) can be used to estimate the level of resistance in *K. scoparia* based on *EPSPS* gene copy number:

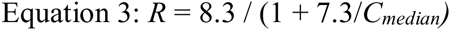

where R is the R:S ratio (or the *n*-fold resistance level), and *C*_*median*_ is the median EPSPS:ALS ratio (or number of *EPSPS* gene copies) in the resistant biotype. If validated independently, this equation could prove useful as it will allow estimation of field resistance level without the need for time-consuming greenhouse bioassays. In laboratories equipped to do this type of work, the *EPSPS* gene copy number can be estimated within a week, whereas a greenhouse bioassay using whole plants can take months, and require either transplanting or growing plants from seed and substantial greenhouse space. Once the *EPSPS* gene copy number is quantified, the level of field resistance could be estimated simply by using Equation 3.

## Discussion

GR *K. scoparia* populations from sugarbeet fields exhibit *EPSPS* gene duplication similar to what is observed in *K. scoparia* from dryland fallow systems. Gene duplication in the tested GR samples has not risen to levels higher than 12 additional copies. Determining the *EPSPS* copy number is a valuable assay for diagnosing glyphosate resistance in *K. scoparia*. If a sample has increased *EPSPS* copy number, our results suggest that the sample is GR (Figs 1 and 3). If a sample does not have increased *EPSPS* copy number, it is glyphosate-susceptible.

Our overall survey results indicate that *EPSPS* copy number in GR *K. scoparia* from sugarbeet fields is generally within the previously observed range of 3-10. *Kochia scoparia* accessions with *EPSPS* copies <3 did not exhibit a field level of glyphosate resistance. An *EPSPS* copy number of 3 would provide approximately 2.4-fold level of glyphosate resistance based on Equation 3, which may be difficult to observe under field conditions. Some accessions had LD_50_ estimates with high standard errors and these were excluded from the analysis. We interpret this as being due to experimental variation in the greenhouse bioassay, and partially due to heterogeneity within the accessions for *EPSPS* copy number. This variance in LD_50_ estimate is not due to somatic instability or loss of *EPSPS* gene duplication in *K. scoparia,* as the inheritance of *EPSPS* gene duplication has been previously shown to be stable [20]. Our analysis here is somewhat limited, since our highest median *EPSPS* copy number for any accession included in the model was 8. Additional *K. scoparia* accessions from eastern Colorado and Alberta, Canada with *EPSPS* copy numbers from 15 to 20 have been recently identified [27]. These populations appear to be highly resistant to glyphosate. Our hypothesis regarding maximum resistance level from this mechanism should be tested empirically with *K. scoparia* populations with higher *EPSPS* copy numbers.

These trends suggest that continued glyphosate selection pressure is selecting for higher *EPSPS* copy number, higher resistance levels, and multiple herbicide resistance in *K. scoparia*. Recently a *K. scoparia* population from Kansas has been confirmed to be resistant to four herbicide modes of action (PSII, ALS, glyphosate, and synthetic auxins) [10]. Known glyphosate resistance mechanisms exceed those reported for any other herbicide and include target-site mutations, target-site gene duplications, active vacuole sequestration, limited cellular uptake, and rapid necrosis response [28]. Proper stewardship of glyphosate is critical, including use of other herbicide modes of action, cultural and mechanical control practices, and preventing seed set on surviving *K. scoparia*.

Our approach is different from many previous calculations of R:S ratio since we included many *K. scoparia* accessions that exhibited a susceptible response. Typically, R:S ratios are calculated using one or two ‘known susceptible’ biotype(s). There is little agreement in the weed science literature on what should be used as a susceptible biotype for resistance confirmation studies [29], and whole-plant response to glyphosate can vary widely among susceptible accessions [30]. Higher *EPSPS* gene copy number increased the estimated probability of survival in GR *Amaranthus palmeri*, when an empirical model fit to an F_2_ population segregating for *EPSPS* gene copy estimated that 53 *EPSPS* gene copies provided a 95% probability of surviving a high dose of 2000 g ae ha^-1^ glyphosate [31]. We used a modeling approach to estimate the level of resistance expected for a single *EPSPS* copy individual, and therefore, can estimate resistance directly attributable to the resistance mechanism. This method of calculating the R:S ratio should be less affected by variability in phenotypic response as a result of different genetic backgrounds. In our study, glyphosate susceptible *K. scoparia* (median *EPSPS* copies <1.5) exhibited glyphosate LD_50_ ranging from 202 to 1225 g ae ha^-1^, indicating glyphosate sensitivity can vary widely among accessions not expressing the resistance mechanism. Similar levels of variability would be expected among plants with multiple *EPSPS* copies. If a single *K. scoparia* accession were used as our ‘known susceptible’ biotype, the R:S ratio of our most resistant accession (LD_50_ = 3895) could range from 3 to 19. Using our empirical model, we provide an estimate of the level of glyphosate resistance that is attributable to the resistance mechanism and is less affected by variability contributed by other, unrelated genetic factors.

## Acknowledgements

This authors thank the Western Sugar Cooperative agronomists for collecting the *K. scoparia* seed samples.

## Supporting Information

**S1 Fig. Greenhouse bioassay results for all *K. scoparia* collections.** A) Glyphosate LD_50_ estimates for the 65 *K. scoparia* accessions included in the bioassay, and B) results for 53 *K. scoparia* accessions after 12 were removed from the analysis due to large standard errors associated with the LD_50_ estimates.

**S2 Fig. Glyphosate resistance level (A) and *EPSPS* gene copy numbers (B) for all 40 *K. scoparia* accessions measured for *EPSPS* gene copy number**. Filled circles represent mean values for each accession; bars represent standard error of the mean; open circles represent median *EPSPS* gene copy number for each accession.

**S3 Fig. Glyphosate resistance level as influenced by *EPSPS* gene copy number in *Kochia scoparia***. Black dashed line represents the fitted model when only filled circles were used in the fitting procedure. Open circles represent *K. scoparia* accessions that were excluded from the model-fitting due to LD_50_ estimates having large standard errors. The blue dotted line and equation represent the fitted model if these points were included in the model fitting procedure.

